# Computational assessment of the spike protein antigenicity reveals diversity in B cell epitopes but stability in T cell epitopes across SARS-CoV-2 variants

**DOI:** 10.1101/2021.03.25.437035

**Authors:** Anni Ge, Melissa Rioux, Alyson A. Kelvin

**Affiliations:** Department of Microbiology and Immunology, Faculty of Medicine, Dalhousie University, Halifax, NS B3H 4R2, Canada; Vaccine and Infectious Disease Organization (VIDO), University of Saskatchewan, Saskatoon, Saskatchewan, S7N-5E3, Canada; Department of Pediatrics, Division of Infectious Disease, Faculty of Medicine, Dalhousie University, Halifax, NS B3K 6R8, Canada; The Canadian Center for Vaccinology (IWK Health Centre, Dalhousie University and the Nova Scotia Health Authority), Halifax, NS B3K 6R8, Canada

## Abstract

Since its emergence into the human population at the end of 2019, SARS-CoV-2 has caused significant morbidity and mortality worldwide. Efforts to develop a protective vaccine against COVID-19 have yielded several vaccine platforms currently in distribution targeting the original SARS-CoV-2 spike protein sequence from the first cases of infection. In recent months, variants of SARS-CoV-2 have raised concerns that viral mutation may undermine vaccination efforts through viral escape of host immune memory acquired from infection or vaccination. We therefore used a computational approach to predict changes in spike protein antigenicity with respect to host B cell and CD8+ T cell immunity across six SARS-CoV-2 variants (D614G, B.1.1.7, B.1.351, P.1, B.1.429, and mink-related). Our epitope analysis using DiscoTope suggests possible changes in B cell epitopes in the S1 region of the spike protein across variants, in particular the B.1.1.7 and B.1.351 lineages, which may influence immunodominance. Additionally, we show that high-affinity MHC-I-binding peptides and glycosylation sites on the spike protein appear consistent between variants with the exception of an extra glycosylation site in the P.1 variant. Together, these analyses suggests T cell vaccine strategies have the most longevity before reformulation.

## 1. Introduction

Severe acute respiratory syndrome coronavirus 2 (SARS-CoV-2) is the highly contagious respiratory virus responsible for the COVID-19 pandemic. As of March of 2021, 118 million COVID-19 cases have been reported, and over 2.36 million deaths have been attributed to the virus worldwide [1]. Host infection with SARS-CoV-2 is initiated by a transmembrane, homotrimeric fusion protein referred to as the spike protein [2]. Vaccination efforts have greatly revolved around the spike protein as an immunogenic target [3,4], as antibody binding to spike effectively blocks viral entry into host cells and suppresses viral infection [5]. Some spike-based vaccine candidates are already in distribution, such as the Pfizer and Moderna mRNA vaccines, as well as the adenovirus vector vaccines from Johnson and Johnson and AstraZeneca [6,7]. Both vaccine platforms have been shown to elicit B cell and T cell specific immunity [8,9].

In recent months, several variants of SARS-CoV-2 have emerged that have raised concerns about the effectiveness of current SARS-CoV-2 vaccine platforms [10]. Initially, the first SARS-CoV-2 variant containing a D614G mutation, which has been shown to stabilize the protein trimer [11], appeared early in the pandemic and rapidly became the dominant circulating virus [12]. In December of 2020, a lineage referred to as B.1.1.7, which had been in circulation in the UK since September, was reported by Public Health England [13]. This variant, modeled to have an increased reproduction number [14], has 17 mutations (14 non-synonymous point mutations and three deletions). Eight of the mutations are contained within the spike protein and include the D614G, a N501Y mutation in the receptor-binding domain of spike; a deletion at position 69-70; and a P681H mutation near the S1/S2 cleavage site [15]. Another lineage designated B.1.351, which has eight mutations within the spike protein including the lineage defining mutations K417N, E484K, and N501Y, was detected in South Africa as early October of 2020 [16,17]. In particular, this observation represents the first instance of the E484K mutation in a variant of concern, which is thought to impact the binding of neutralizing antibodies [18]. A third significant variant of concern, referred to as P.1 from the B.1.1.28 lineage, was identified out of Brazil in late 2020 with several mutations similar to those of the B.1.351 mutant, suggesting convergent evolution [19]. Sequencing found 17 amino acid changes, three deletions, and four synonymous mutations in the protein with one four-nucleotide insertion. Other variants of importance include the B.1.429 [20] identified in California and the mink-related lineages detected in Denmark in June of 2020, which also have several non-synonymous mutations compared to the Wuhan SARS-CoV-2 reference sequence [21]. The mink variants have been suggested as potential future spillover viruses, since minks may provide a viral reservoir that has frequent interaction with people. However, at this time, spread of the mink-related variant to humans has been rare [22].

Mutations across SARS-CoV-2 variants are predicted to affect several key functions of the virus, including strengthening viral binding to the host angiotensin converting enzyme 2 (ACE2) [23], increasing viral infectivity [24], and impacting viral susceptibility to neutralization by host immune responses [12]. Through several lines of investigation such as *in vitro* neutralization assays, *in vivo* challenge studies, and human epidemiological studies, it is important to determine the protective effectiveness afforded by vaccination or previous infection against these variants. At this time, it is still unclear whether SARS-CoV-2 variants of concern are able to be inhibited by previously acquired host adaptive immune response. Furthermore, as vaccine effectiveness is dependent on the recognition of viral proteins by T and B cells, mutations influencing T and B cell epitopes in new SARS-CoV-2 variants may have significant effects on the performance of vaccines developed toward the original Wuhan SARS-CoV-2 strains. Therefore, to gain a molecular understanding of vaccine performance against the new variants, we assessed the antigenicity of SARS-CoV-2 spike protein of the current variants of concern including the D614G, B.1.1.7, B.1.351, B.1.429, P.1, and mink-related mutants. Using complete spike sequences obtained from the Global Initiative on Sharing All Influenza Data (GISAID) [25], we applied rationally chosen computational approaches to compare B cell epitopes, peptides presented to host CD8+ T cells, and glycosylation sites across SARS-CoV-2 variants. We report that some variants of concern have differences in predicted B cell epitopes within the S1 subunit of the spike protein. However, with respect to peptides presented on human MHC-I molecules, the top T cell epitope predictions appear to be consistent between spike variants, suggesting stabilized immunodominance. With this data we conclude that T cell vaccine strategies may have the broader coverage compared to B cell directed strategies against circulating variants and the greatest longevity before vaccine reformulation is required.

## 2. Materials and Methods

### Alignment of variant SARS-CoV-2 sequences

Amino acid sequences for the original Wuhan SARS-CoV-2 spike protein, as well as the SARS-CoV-2 variants D146G, B.1.1.7, B.1.351, B.1.429, P.1, and mink-related SARS-CoV-2, were obtained from GISAID [25]. The accession numbers for each amino acid sequence are as follows: Wuhan SARS-CoV-2 (EPI_ISL_402124), D614G (EPI_ISL_406862), B.1.1.7 (EPI_ISL_852526), B.1.351 (EPI_ISL_864621), B.1.429 (EPI_ISL_1017160), P.1 (EPI_ISL_1171653), and mink-related (EPI_ISL_615652). We compared variant-specific mutations across the various spike proteins by aligning amino acid sequences in MEGAX (Molecular Evolutionary Genetics Analysis, Version 0.1) [26], with deletions denoted by dashes. Alignment figures were generated in ClustalX Version 2.0 [27].

### Modelling spike protein mutations

To predict the three-dimensional structure of each variant protein, we applied the web-based bioinformatics server Phyre2 (Protein Homology/ analogY Recognition Engine Version 2.0 (http://www.sbg.bio.ic.ac.uk/~phyre2/html/page.cgi?id=index)) [28]. Intensive modelling mode was selected for the amino acid sequence of each variant. The templates applied in modelling folding of reference and variant spike sequences are as follows: c7dk3B, c6vsbB, c2kncA, c7a4nA, c6vsbA, c2fxpA, c3j3bG, c6vybB, c6zowB, c1m0jA, c1s6wA, c5h36E (see **Table S1**). The model coordinate files in Protein Database (PDB) format were downloaded for downstream visualization and manipulation. We created structural images of each variant with PyMOL (The PyMOL Molecular Graphics System, Version 2.3.5, Schrödinger, LLC). Maximum performance quality was selected, and proteins were viewed as “sphere” models with the variant-defining mutations highlighted. Next, the web-based software Missense3D (http://www.sbg.bio.ic.ac.uk/~missense3d/) [29] was employed to predict the structural impact of each individual variant amino acid substitution. For this analysis, the wild-type spike sequence was set as the reference, and substitutions across all six SARS-CoV-2 variants were input manually. A cavity volume expansion or contraction of ≥ 70 Å^3^ was defined as structural damage to differentiate from minor expansions or contractions.

### Prediction of B cell epitopes across variants

Predicted B cell epitopes across spike variants were assessed using the DiscoTope 2.0 server (http://www.cbs.dtu.dk/services/DiscoTope/) [30]. The corresponding PDB file of each variant was input at three different threshold scores for defining positive B cell epitopes: the least conservative score of −3.7 (0.47 sensitivity, 0.75 specificity), a moderately conservative score of −2.5 (0.39 sensitivity, 0.80 specificity), and the most conservative score of +1.9 (0.17 sensitivity, 0.95 specificity). The DiscoTope output was used to generate heatmaps of B cell epitope probability across variant protein residues in PyMOL, with predicted epitopes in yellow. The amino acid number and DiscoTope score were used to map variant B cell epitopes in RStudio (version 3.6.1) using the ggplot2 package (version 3.3.2)[31].

### Assessment of predicted T cell peptides

The likelihood of SARS-CoV-2 peptides being presented on human MHC-I molecules was predicted using the Immune Epitope Database (IEDB) Analysis Resource NetMHCpan EL 4.1 (http://tools.immuneepitope.org/main/). The complete amino acid sequences for the Wuhan and variant SARS-CoV-2 spike proteins were used, and predicted peptides were restricted to 27 common human HLA genes to encompass most of the human population [32,33]. The T cell epitopes predicted for the Wuhan and variant SARS-CoV-2 were compared by filtering the data in RStudio using the tidyverse package (version 1.3.0) [34].

### Variant spike glycosylation site analysis

Wild-type and variant amino acid sequences were uploaded to the NetNGlyc 1.0 server (http://www.cbs.dtu.dk/services/NetNGlyc/) [35] to assess potential N-linked glycosylation sites (Asn-Xaa-Ser/Thr sequons). The default threshold of 0.5 was used. The predicted glycosylation sites across variants were modelled using PyMOL (The PyMOL Molecular Graphics System, Version 2.3.5, Schrödinger, LLC).

## 3. Results

Six distinct SARS-CoV-2 variants of concern, including D614G, B.1.1.7, B.1.351, B.1.429, P.1, and the mink-related variant, were selected along with the original SARS-CoV-2 Wuhan virus (herein referred to as the “reference” spike) to explore antigenic differences in response to mutations within their surface glycoprotein. We first aligned reference and variant spike sequences using the MEGAX alignment tool [26] to better visualize the relative location of sequence mutations (**Figure 1**). Across variants, twentyseven substitutions (S13I, L18F, T20N, P26S, D80A, W152C, D138Y, R190S, D215G, K417T, K417N, L452R, Y453F, E484K, N501Y, A570D, D614G, H655Y, P681H, I692V, A701V, T716I, S982A, T1027I, D1118H, V1176F, and M1229I) and deletions at position 69-70, 145, and 241-243 were noted (**Figure 1A**). A common D614G mutation was identified in all six variants, and the substitution at position 501 within the receptor-binding domain was shared among B.1.1.7, B.1.351, and P.1 lineages (**Figure 1B**). Moreover, the mink-related variant contained a deletion that was also recognized in B.1.1.7 at positions 69-70.

**Figure 1.**
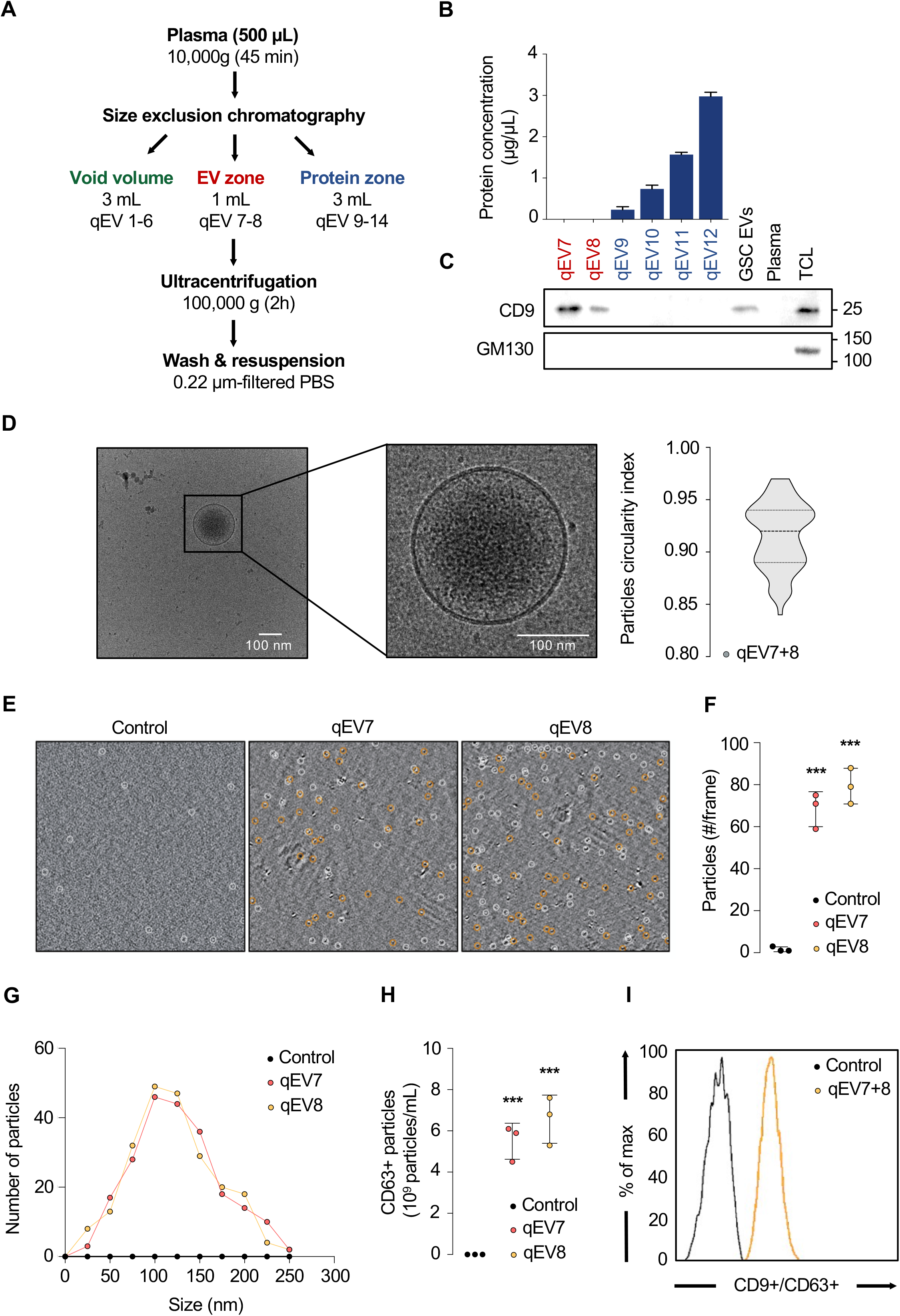
Amino acid sequence changes in variant and Wuhan SARS-CoV-2 spike protein. Sequences for Wuhan (reference) SARS-CoV-2 spike, as well as from variants D614G, B.1.1.7, B.1.351, B.1.429, P.1, and mink-related SARS-CoV-2, were obtained from the GISAID [6] and are as follows: the original Wuhan SARS-CoV-2 reference sequence (EPI_ISL_402124); B.1.1.7 (EPI_ISL_852526); B.1.351 (EPI_ISL_864621); B.1.429 (EPI_ISL_1017160); D614G lineage (EPI_ISL_406862), P.1 (EPI_ISL_1171653), and the mink-related lineage (EPI_ISL_615652). **(A)** Complete spike sequences were aligned using MEGAX. Differences between sequences are indicated with arrows, and deletions are indicated with dashes. The figure was generated using the ClustalX platform. **(B)** Three-dimensional models of variant spike monomers highlighting detected amino acid substitutions were produced using PyMOL (The PyMOL Molecular Graphics System, Version 2.3.5, Schrödinger, LLC).

It is unclear how the amino acid substitutions or deletions across variants will impact SARS-CoV-2 spike structure and function. We aimed to characterize the structural impact of each mutation using the web-based server Missense 3D [29] (**Table S2**). Using the Wuhan SARS-CoV-2 spike as a reference, the individual amino acid substitutions were assessed for structural changes such as buried charged residues, disruption of salt bridges, and major cavity expansions or contractions. Of the twenty-seven substitutions spanning the six spike variants, notable structural damage was only predicted for four mutations. First, the aspartate to alanine switch at residue 80 (D80A) in the B.1.351 variant was predicted to change a buried, charged residue to an uncharged residue and disrupt the corresponding salt bridge. Second was the proline to histidine switch at residue 681 (P681H) in the B.1.1.7 spike protein, which was predicted to cause a switch from an uncharged, buried amino acid residue to an exposed, positively charged residue. Finally, two mutations exclusive to the P.1 variant revealed structural damage: L18F was predicted to contract the cavity volume by 104.976 Å^3^, while D138Y contracted the cavity volume by 91.584 Å^3^ and disrupted the buried H-binding, causing a buried amino acid to be switched to an exposed residue. The remaining amino acid substitutions were not predicted to cause structural changes. Because Missense 3D evaluates missense mutations, the deletions appearing in B.1.1.7, B.1.351, and mink-related spike were not assessed in this particular analysis.

### 3.1. Predicted B cell epitopes differ across SARS-CoV-2 spike protein variants

Although predicted changes in SARS-CoV-2 spike structure can help elucidate receptor docking as well as protein function such as membrane fusion, our interests were in understanding the immune system recognition of the viral variants and the stability of antigenicity across variants. Therefore, the primary goal of our analysis was to evaluate changes in spike antigenicity. To this end, we first aimed to compare the predicted B cell epitopes across reference and variant spike proteins to determine whether the variants may present differently to host B cell receptors and thereby have differential recognition by secreted antibodies, which may have been elicited from previous exposures. For this analysis, we used scores generated by the DiscoTope 2.0 server from DTU Heath Tech to identify epitopes of interests and determine differences between the variants [30]. We selected three DiscoTope score thresholds with differing levels of sensitivity and specificity: first, the default −3.7-score threshold which is least conservative when selecting B cell epitopes (**Figure 2**), then a moderately conservative threshold of −2.5, followed by a +1.9 threshold, the most conservative (**Table S3**). In examining the positively identified epitopes at the −3.7 threshold, we found that the number of residues predicted to be involved in B cell epitopes differed across the variants. Overall, 168, 166, 154, 172, 143, 145, and 149 residues were predicted as positive B cell epitopes in the Wuhan reference, B.1.1.7, B.1.351, B.1.429, D614G, P.1, and the mink-related spike protein sequences, respectively. We found that changes to specific B cell epitopes occurred almost exclusively within the S1 subunit and protease cleavage regions of the spike protein in all variants. In plotting the DiscoTope scores across the amino acid number of each variant, we noted specific differences in regions predicted to interact with host B cells. In the D614G, B.1.1.7, and P.1 variants, a high-scoring epitope was identified spanning residues 677-688 in the SD1/2 region of spike, which is part of the S1 subunit [36] (**Figure 2B, 2C**). Another epitope at residues 244-250 detected in the reference and other variants was absent in B.1.351, suggesting this particular epitope may be less accessible to host humoral responses compared to the other variants (**Figure 2B**). We also noted an epitope at residue 180 that scored lower in the B.1.429 and mink-related lineages (**Figure 2B**). Taken together, these results demonstrate that the specific amino acid substitutions in the S1 subunit of spike may lead to differential recognition or evasion of SARS-CoV-2 variants by specific B cell clonotypes elicited after infection.

**Figure 2.**
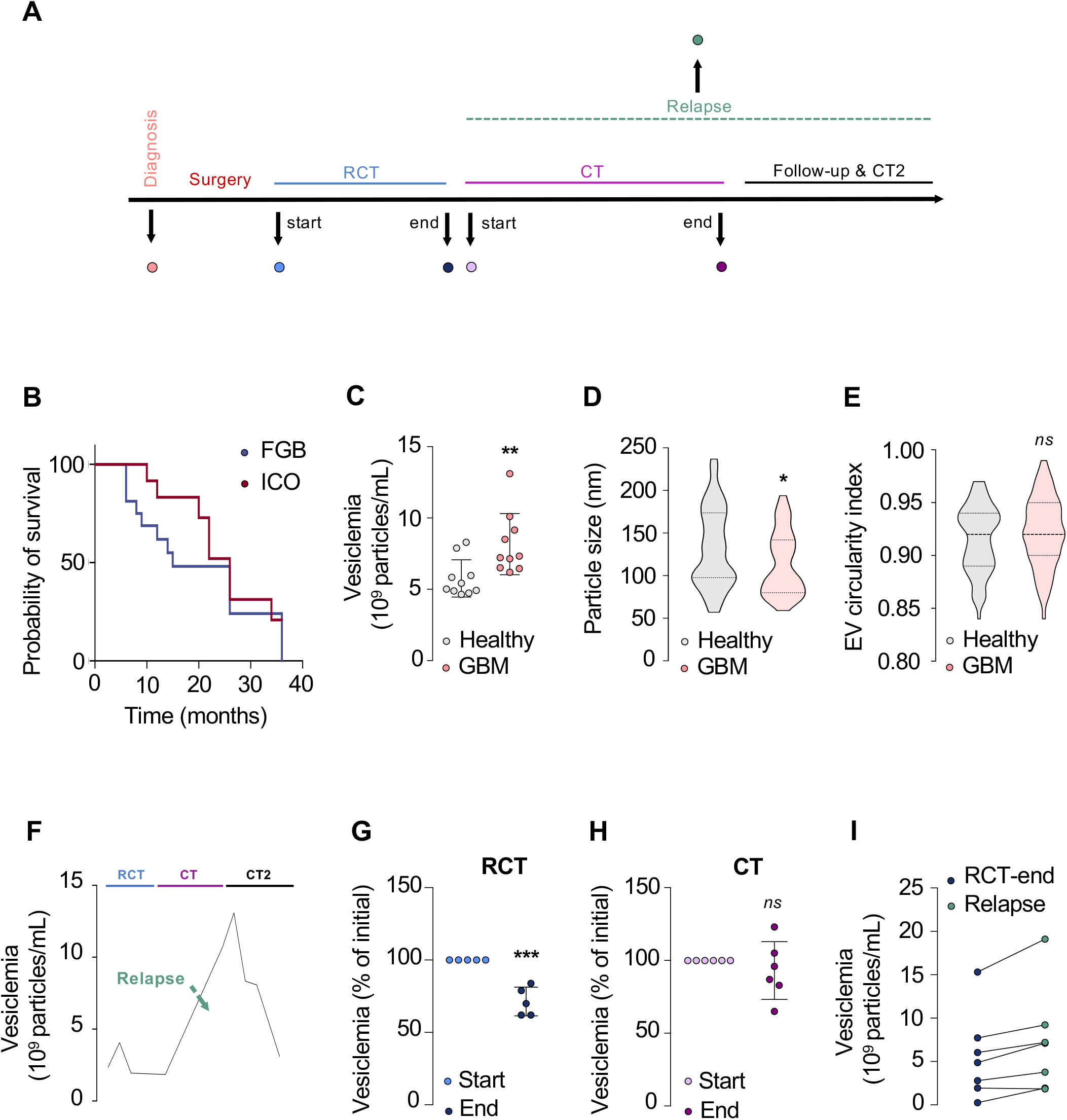
Changes in predicted B cell epitopes in spike proteins of SARS-CoV-2 variants compared to the Wuhan SARS-CoV-2 sequence. Antigenic analysis of the spike protein of SARS-CoV-2 variants was performed using the DiscoTope 2.0 platform (DTU Health Tech) (http://www.cbs.dtu.dk/services/DiscoTope/_)[11]. Sequences were obtained from the GISAID database [6] and are as follows: the original Wuhan SARS-CoV-2 reference sequence (EPI_ISL_402124); B.1.1.7 (EPI_ISL_852526);B.1.351(EPI_ISL_864621);B.1.429(EPI_ISL_1017160);D614G lineage (EPI_ISL_406862), P.1 (EPI_ISL_1171653), and the mink-related lineage (EPI_ISL_615652). **(A)** DiscoTope scores falling below (blue) and above (red) the B cell epitope prediction threshold of −3.7 (0.47 sensitivity, 0.75 specificity) were mapped along the amino acid sequences of reference (Wuhan) and variant (D614G, B.1.1.7, B.1.351, B.1.429, P.1, and mink-related) SARS-CoV-2 spike proteins. **(B)** Only the positive B cell epitope prediction results are shown for greater resolution of the B cell epitope regions. The arrows highlight differences in predicted epitopes within the variants of concern compared to the reference spike protein. **(C)** Spike monomers with DiscoTope scores represented as a heatmaps were mapped along the protein. Predicted epitopes for each group are highlighted in yellow, and blue arrows denote regions that are structurally/antigenically different. The image was generated using PyMOL (The PyMOL Molecular Graphics System, Version 2.3.5, Schrödinger, LLC).

### 3.2. High-affinity MHC-I-binding peptides conserved across variants

Along with altering possible B cell epitopes, mutations in amino acid sequences can lead to changes in antigenicity with respect to the specific peptides presented to host T cells. Importantly, the T cell epitopes (CD8+ T cell epitopes binding MHC-I) are linear, consisting of approximately 8-11 amino acids, while B cell epitopes typically require a 3D confirmation of discontinuous amino acids for recognition [37]. Since both arms of the immune system are important during antigen encounter, we next assessed the amino acid substitutions and deletions across SARS-CoV-2 variants for changes in T cell epitopes that may differentially stimulate host CD8+ T cells. Using the complete amino acid sequences for reference and variant spike, we generated MHC-I binding predictions using the Immune Epitope Database (IEDB) Analysis Resource NetMHCpan EL 4.1 {Citation}. We restricted our analysis to 27 HLA-A and HLA-B alleles to encompass most of the human population [33]. In total, we found variants to differ in the number of peptides predicted to bind to host MHC-I molecules: reference, D614G, and B.1.429 each had 68,283 unique HLA allele-peptide combinations, while mink-related spike had 68,175; B.1.1.7 had 68,121; P.1 had fewer at 52,899 epitopes; and B.1.351 had 51,569 epitopes. We filtered the data such that peptides with the highest MHC-I binding affinity were selected for each group (percentile rank < 1). Here we found that variants differed in the predicted number of high-affinity peptides that can be presented across human MHC-I alleles: the reference, D614G, and mink-related spike each had 597 peptides, while B.1.1.7 had 595, B.1.429 had 594, and P.1 and B.1.351 had additional peptides, at 604 and 602, respectively. Although there were differences in total numbers of high-affinity peptides among the variants, the overall top 20 high-scoring peptides were conserved among the reference and variants. The top 20 conserved peptides between reference and variant sequences are indicated (**Table 1**). However, we found that all variants containing the D614G mutation lacked three high-affinity peptides present in the reference spike protein (**Table 1**). These are DVNCTEVPV (614-622) and QVAVLYQDV (607-615) on the HLA-A*68:02 allele, and YQDVNCTEV (612-620) on alleles HLA-A*02:06 and HLA-A*02:01 (**Table 1**). Additionally, our analysis detected several unique MHC-I-binding peptides per variant that were not present in the reference sequence or other variants (**Table S4**). The B.1.1.7 lineage had 25 unique peptides, B.1.351 had 24, B.1.429 had 15, P.1. had 37, and mink-related spike had 20. These peptides spanned both S1 and S2 subunits of spike. Of the 25 high-affinity peptides specific to the B.1.1.7 lineage, 11 were in the S1 subunit, but none spanned the receptor binding domain of spike (**Table S4**). This result was similar to the mink-related lineage, which had approximately half its variant-specific peptides occurring in S1 but not in the RBD, as well as B.1.429, which had all unique peptides in S1 but none in RBD. The P.1 lineage had the most variant-specific peptides, with 27 of its unique peptides occurring in S1, and seven falling in the RBD specifically (**Table S4**). With respect to CD8+ T cell activation, the B.1.1.7, B.1.429, and mink-related variants appear antigenically similar to reference SARS-CoV-2 across the RBD of spike. However, P.1 and B.1.351 lineages show RBDs that may have antigenic distinctions from the reference. With the exception of B.1.1.7, we found the MHC-I binding peptides derived from the S2 subunit of spike to be more conserved between variant and reference spike proteins.

**Table 1.**
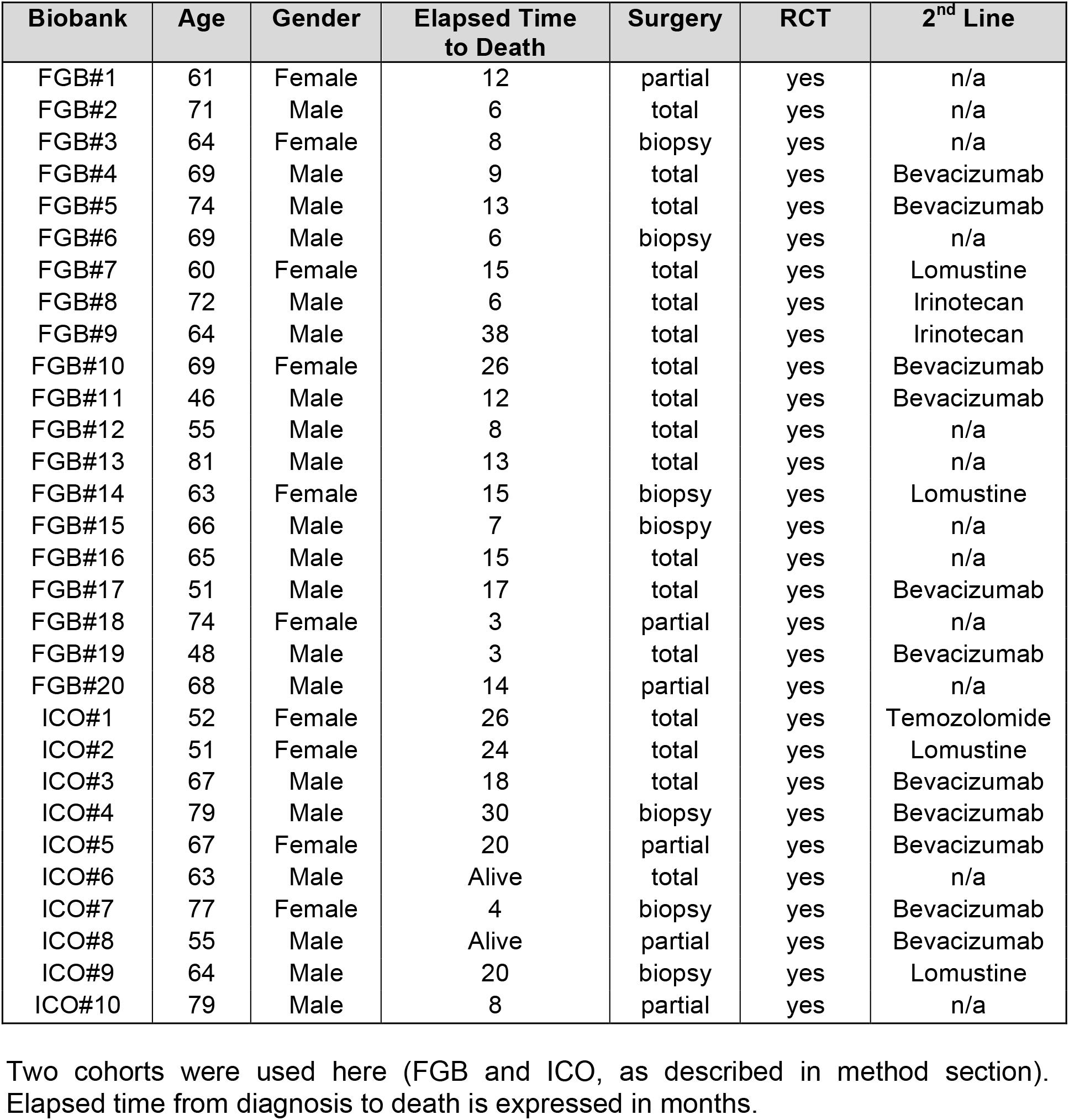
Top 20 high-affinity MHC-I-binding peptides predicted by NetMHCpan EL 4.1 (http://tools.immuneepitope.org/main/) consistent across reference SARS-CoV-2 and D614G, B.1.1.7,B.1.351, B.1.429, P.1, and mink-related variants, as well spike peptides lost in variant SARS-CoV-2 spike.

Conservation of the top spike peptides across human HLA alleles among the reference and variant SARS-CoV-2 viruses suggests that cellular adaptive immune responses may still be effective despite the mutations. However, our analysis also indicated that lineage-defining mutations span sequences likely to be presented on common human HLA alleles, possibly leading to some variant-specific CD8+ T cell immunity following exposure. Immunodominance will be an important next step in the analysis of the adaptive immune response to SARS-CoV-2 variants.

### 3.3. Glycosylation site predictions are consistent across SARS-CoV-2 variants

Antigen glycosylation is an important factor in host immunity to pathogens. Glycosylation of viral envelope proteins can modify the protein such that it can increase infectivity [38] and shield it from host immune response [39–41]. The spike protein of SARS-CoV-2 is known to be heavily glycosylated [42,43]. To assess whether spike mutations impact the predicted N-linked glycosylation sites (N-X-S/T, X ≠P) in SARS-CoV-2 variants, and thus possibly antigenicity, we used the NetNGlyc 1.0 server from DTU Health Tech (http://www.cbs.dtu.dk/services/NetNGlyc/) [35] using the default threshold of 0.5. The data derived from the server suggested that seventeen conserved locations that crossed the threshold were found in all variants with similar scores at sites 17, 61, 74, 122, 149, 165, 234, 282, 331, 343, 603, 616, 717, 801, 1098, 1134, and 1194 (**Table S5**). Three additional sites were detected in the P.1 variant with only (+) N-glycosylation results at positions 20, 188, and 657. However, only 7 sites (61, 74, 234, 282, 616, 717, and 1194) with (++) or better predictions were selected to generate the 3D structure of spike protein with glycans for high specificity (**Figure 3**), and the P.1 variant was found to contain one more glycosylated site at position 17 with (++) score than the rest of the strains, which shared the same 7 N-linked sequons. No differences were observed between the other five variants of concern, including D614G, B.1.1.7, B.1.351, B.1.429, and the mink-related spike, whereas the mutations present exclusively in the P.1 variant could alter the glycosylation process.

**Figure 3.**
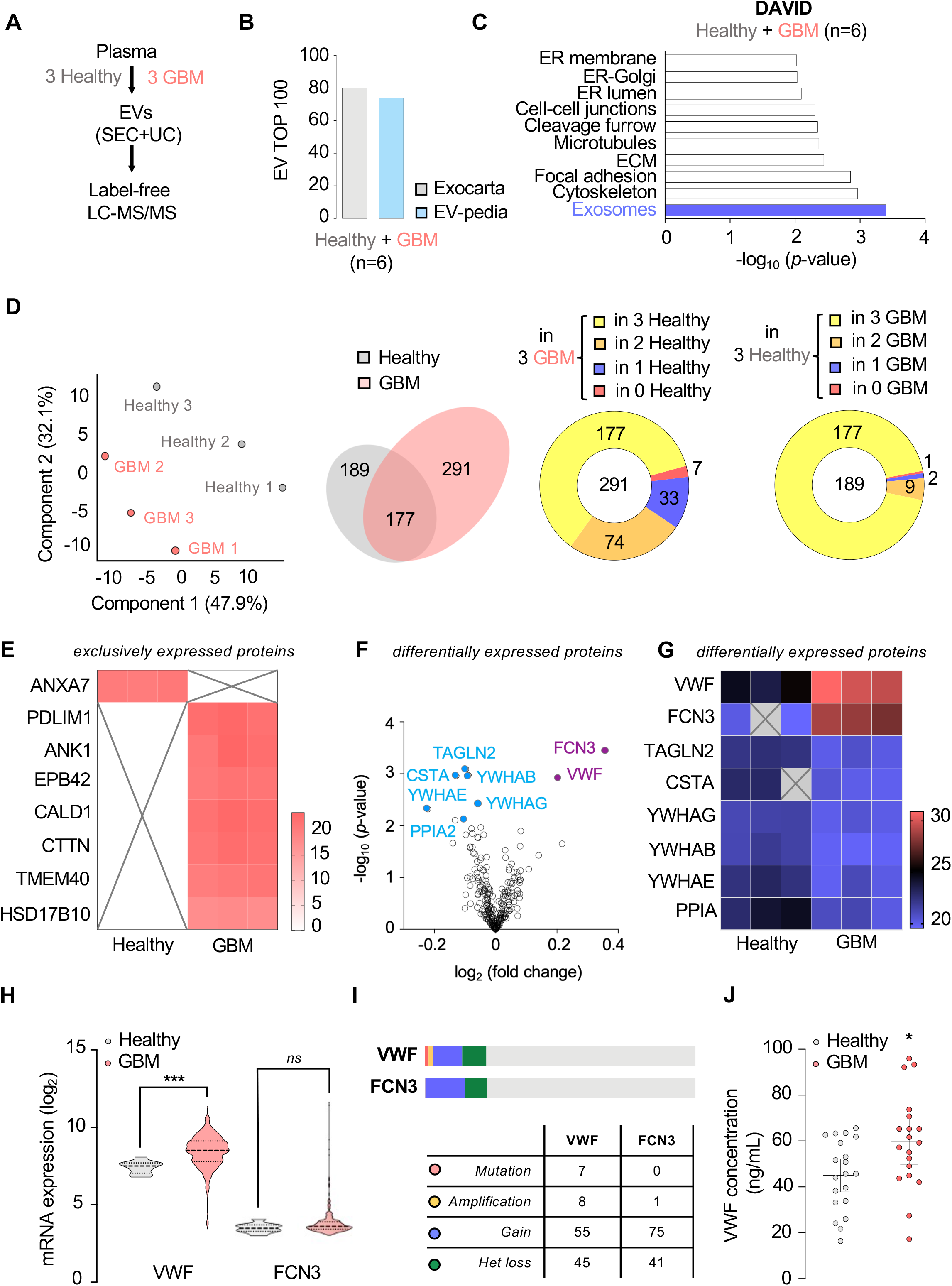
P1 variant has an additional high-scoring glycosylation site. N-glycosylation sites on reference and variant SARS-CoV-2 amino acid sequences were predicted using the NetNGlyc 1.0 server. Predicted sites were filtered to show only those with (++) and (+++) NetNGlyc scores, revealing seven sites. These sites are predicted to be shared between D614G, B.1.1.7, B.1.351, B.1.429, P.1, and mink-related spike proteins. The figure shows a representative spike monomer with glycans in green at the predicted sites. The model was generated in PyMOL (The PyMOL Molecular Graphics System, Version 2.3.5, Schrödinger, LLC).*Site 8 indicates an additional high-scoring glycosylation site in the P.1 variant only.

## 4. Discussion

Here we investigated the antigenic changes in the SARS-CoV-2 spike protein among the original viral sequence and the newly identified variants of concern. Antigenicity of a protein is essential for eliciting a protective immune response. The classic evolutionary arms race between virus and host often leads to the virus gaining mutations that allow evasion from previously acquired adaptive immunity. We found that the six SARS-CoV-2 variants of concern in our investigation had minor, yet notable, changes in antigenicity from the amino acid sequence changes. Notable changes were identified in the B.1.351 and the P.1 variants, which may account for the decreases in neutralizing ability of convalescent and vaccine serum against new variants, as the majority of the B cell antigenic changes were identified in the receptor binding domain. Additionally, we determined that the top 20 MHC-I binding predictions were conserved across the variants. These data are important for assessing the performance of current COVID-19 vaccines against changes in the SARS-COV-2 virus, considering new vaccine formulations, and developing nextgeneration vaccine platforms.

The spike protein is the primary target for existing SARS-CoV-2 vaccine platforms. Because it is key for viral entry and expressed on the virion surface, spike is recognized by either antigen-presenting cells or T cells, which are essential for production of antigen-specific neutralizing antibodies [44]. Therefore, structural changes in the variant spike proteins could influence recognition by immune cells which is required to elicit appropriate cellular and humoral immune mechanisms for clearing virus. No potential structural damage or major changes were detected with the exception of four substitutions found in B.1.1.7 (P681H), B.1.351 (D80A), and P.1 (L 18F and D138Y). Some of the identified substitutions have already been functionally investigated and found to affect the transmissibility and antigenicity of the virus. Specifically, N501Y present in the receptor-binding domain (RBD), which is essential for viral entry, has an increased angiotensin converting enzyme 2 (ACE2) receptor binding affinity with or without the help of the deletion at 69-70 [43]. Additionally, SARS-CoV-2 containing the E484K substitution showed a reduction in antibody neutralization ability in several studies [18], and the L452 mutation was said to confer resistance to binding by certain monoclonal antibodies [20,45] along with possibly increasing infectivity [46]. The mutation K417T present in the RBD, along with E484K and N501Y, is described as part of a trio of mutations that likely increase transmissibility of the P.1 variant [19]. The well-studied mutation D614G that quickly became the dominant form early in the pandemic is shown to enhance not only the transmissivity of SARS-CoV-2, but also receptor binding domain-specific antibody susceptibility and trimer stability, although we did not predict a structural effect from Missense3D in our analysis [11,12,24,47]. Other spike mutations are shown to impact viral fitness directly. The L18F mutation, for which we identified structural damage in our analysis of the P.1 variant, is said to confer a replicative advantage [48].

Once the variants were identified, essential studies were initiated to determine if antibodies elicited by vaccination or by previous infection with the original Wuhan SARS-CoV-2 virus could neutralize the new variants. In general, vaccination appears to provide lower neutralizing antibody titers compared to polyclonal antibodies elicited after infection against the variants [49]. Vaccine-induced immunity provides more robust heterotypic immunity than natural infection to emerging SARS-CoV-2 variants of concern [49]. Neutralization assays performed across the variants with vaccinee serum have indicated modest decreases in titers against the B.1.1.7 lineage, with more pronounced losses in titer for the variants which contain the E484K substitution (P.1 and B.1.351) [50–53]. SARS-CoV-2 variant B.1.1.7 is susceptible to neutralizing antibodies elicited by ancestral spike vaccines. In parallel, vaccine clinical trials held in regions with high dominance of a particular variant have demonstrated similar findings, leading to the conclusion that vaccines have greater cross-protective immunity against the B.1.1.7 compared to others, although further analysis is still needed due to small numbers in study participants [54,55]. Our analysis points to additional B cell epitope changes that may account for these results.

Here we found that the major T cell epitopes were conserved across the variants. Our data supports a similar analysis by Tarke and colleagues, which also determined negligible impact of SARS-CoV-2 variants with respect to CD8+ T cell epitopes [56]. Additionally, the bioinformatic analysis was supported by experimental evidence illustrating similar CD4+ and CD8+ T cell simulation by SARS-CoV-2 variant peptide pools. Another study, however, suggests the spike protein of SARS-CoV-2 as a poor CD8+ T cell target since the reactivity was less than that of other spike proteins from the previous coronaviruses SARS-CoV-1 and MERS-CoV (Middle Eastern respiratory syndrome coronavirus) [57]. Despite this analysis, our work, together with the evidence that the current vaccine platforms can elicit strong T cell responses, suggests the importance of developing T cell eliciting vaccines for an RNA virus with a propensity to mutate. More work is needed to isolate the T cell response to determine if T cells alone can provide cross-protection against SARS-CoV-2 infection and COVID-19.

Although we found a broad conservation of strong T cell epitopes and minimal B cell changes in antigenicity, it is important to consider how small-scale antigenic changes in viral proteins can lead to evasion of the host immune system. We noted differences in the predicted B cell epitopes of SARS-CoV-2 variants of concern. In particular, the B.1.351 variant showed loss of a predicted epitope at residue 244 that was present in the reference and other variants investigated in this analysis. Additionally, a high-scoring B cell epitope appears at residue 677 in the B.1.1.7 variant. These changes may affect immunodominance, leading to immune skewing during reinfection or the generation of immunodominant clonotypes, which may be unfavorable in a vaccine.

Host post-translational modification is key for the virus to evade immune responses and mediate protein folding. The SARS-CoV-2 spike gene encodes 22 N-linked glycan sequons per protomer. Overall, 40% of the trimeric S protein surface that contained 66 glycosylated sites was covered by glycans, which can prevent antibody recognition regardless of the carbohydrate types. One study suggested that the ACE2 binding domain that is the most important epitope is unlikely to be affected by antigenic drift for evading immune response without attenuated fitness [43]. The integrity of the RBD might play a crucial role in vaccine design that could maintain its efficiency, in spite of mutations. However, the specific glycans might influence both innate and adaptive immune responses by affecting the spike protein binding through pattern recognition receptors (PRRs), like collectins and other lectins, and via changing the accessible HLA antigens. The receptor binding sites of SARS-CoV-2 are shielded by proximal glycosylated sites (N165, N234, and N343) from recognition of immune cells [43,58], and this is a common feature of viral glycoproteins, which is observed in other viruses (SARS-CoV-1 S, HIV-1 Env, influenza HA, LASV GPC). Furthermore, it has been well documented that influenza viruses that spill over into humans from the animal reservoir have a low level of glycosylation within the HA molecule [59,60]. Interestingly, as the virus becomes more humanized and less like the virus from its original animal host, the virus gains mutations facilitating glycosylation sites [59,60]. The increased glycosylation sites are thought to decrease antibody recognition and neutralization, thereby providing a fitness advantage for the virus. It will be important to continue to monitor the SARS-CoV-2 spike protein for increases in glycosylation sites as it circulates in humans.

## 5. Conclusion

Here we used a bioinformatic approach to analyze the antigenicity of the spike protein sequence among SARS-CoV-2 variants in an effort to gain insight into vaccine performance and inform vaccine design. Our study should be followed up with immune cell cross-reactivity studies, either utilizing convalescent or vaccine sera or in animal models. We identified B cell epitopes in new variants and also determined a relatively stable glycosylation signature with T cell epitope conservation across the variants. Our established pipeline can be used for the analysis of future SARS-CoV-2 variants.

## Acknowledgements

We greatly acknowledge the review of this manuscript by Dr. Darryl Falzarano and Dr. Kristen Kindrachuk. This article is published with the permission of the Director of VIDO-InterVac.

